# A Molecular Dynamics Protocol for Rapid Prediction of EGFR Overactivation and Its Application to the Rare Mutations S768I, S768N, D761N

**DOI:** 10.1101/2025.02.27.640703

**Authors:** Julian Behn, R. N. V. Krishna Deepak, Jiancheng Hu, Hao Fan

**Author notes:** Correspondence to Hao Fan: Bioinformatics Institute (BII), Agency for Science, Technology and Research (A*STAR), Singapore 138671, Republic of Singapore. E-mail addresses and. Declarations of interest: none.

## Abstract

Hyperactivation caused by mutations in the Epidermal Growth Factor Receptor (EGFR) kinase domain is implicated in various diseases, including cancer. However, the structural mechanisms underlying overactivation in many EGFR mutations remain poorly understood, and exploring these mechanisms through conventional experiments or *in silico* simulations is often labor- and cost-intensive. Here, we establish a Molecular Dynamics (MD) protocol capable of rapidly revealing EGFR mutant modes of action using multiple short simulations. We first simulated wild-type EGFR and the well-studied oncogenic mutations L858R and T790M/L858R under different simulation conditions, to derive a protocol which could recapitulate their experimentally established behavior. We then applied this protocol to three rare EGFR mutations: S768I, S768N, and D761N. Experimental studies have suggested that S768I and D761N are oncogenic, whereas S768N is likely a neutral mutation that does not significantly alter EGFR activity. Our simulation results were consistent with these functional indications and provided the corresponding molecular bases – S768I and S768N affect the orientation and stability of the catalytically important αC-helix, while D761N introduces a new hydrogen bonding network between the αC-helix and activation loop. Collectively, the protocol presented here provides a robust and rapid framework for characterizing EGFR mutation mechanisms and is readily adaptable to novel or uncharacterized variants.

## 1. Introduction

The Epidermal Growth Factor Receptor (EGFR) is a transmembrane tyrosine kinase receptor of significant clinical importance. Mutations in its intracellular kinase domain play a pivotal role in cancer development, particularly in Non-Small Cell Lung Cancer (NSCLC), the most prevalent form of lung cancer [1]. The prevalence of EGFR mutations in NSCLC varies demographically, ranging from 10 – 20% in Caucasians to 40 – 60% in Southeast Asians [2]. Targeted therapy using tyrosine kinase inhibitors (TKIs) has significantly improved treatment outcome for EGFR-related cancers [3]. However, treatment failure due to acquired or de novo resistance remains a major challenge in EGFR-targeted therapy [4]. Understanding the structural basis of oncogenic EGFR mutations is therefore essential for advancing therapeutic strategies.

The intracellular kinase domain of EGFR adopts a characteristic kinase fold, with an N-lobe and a C-lobe linked by a short hinge region (Figure 1a). This domain is highly dynamic and transitions between multiple conformations [5]. In the catalytically active conformation, the interlobal activation loop extends outward, allowing substrate access to the ATP pocket. Active kinases are also characterized by an inward-facing αC-helix. This αC-helix orientation facilitates the formation of the crucial K745 – E762 salt bridge, which aids in the coordination of the ATP – Mg^2+^ complex. Additionally, ATP binding is supported by the proper positioning of the highly conserved Asp-Phe-Gly (DFG) motif. In contrast, the ‘Src-like’ inactive conformation features a collapsed activation loop that obstructs the substrate-binding ATP pocket, an outward-facing αC-helix, and a disrupted K745 – E762 salt bridge. A second inactive EGFR conformation, the ‘DFG-out’ conformation, involves the DFG motif blocking ATP binding. Most experimentally resolved EGFR structures, however, fall into either the active or Src-like inactive conformation, while in silico simulations have identified additional partially disordered states that may regulate EGFR activity [6–9]. In the absence of an activating ligand, wild-type EGFR predominantly favors the inactive state, and its conformational dynamics are further influenced by dimerization.

**Figure 1.**
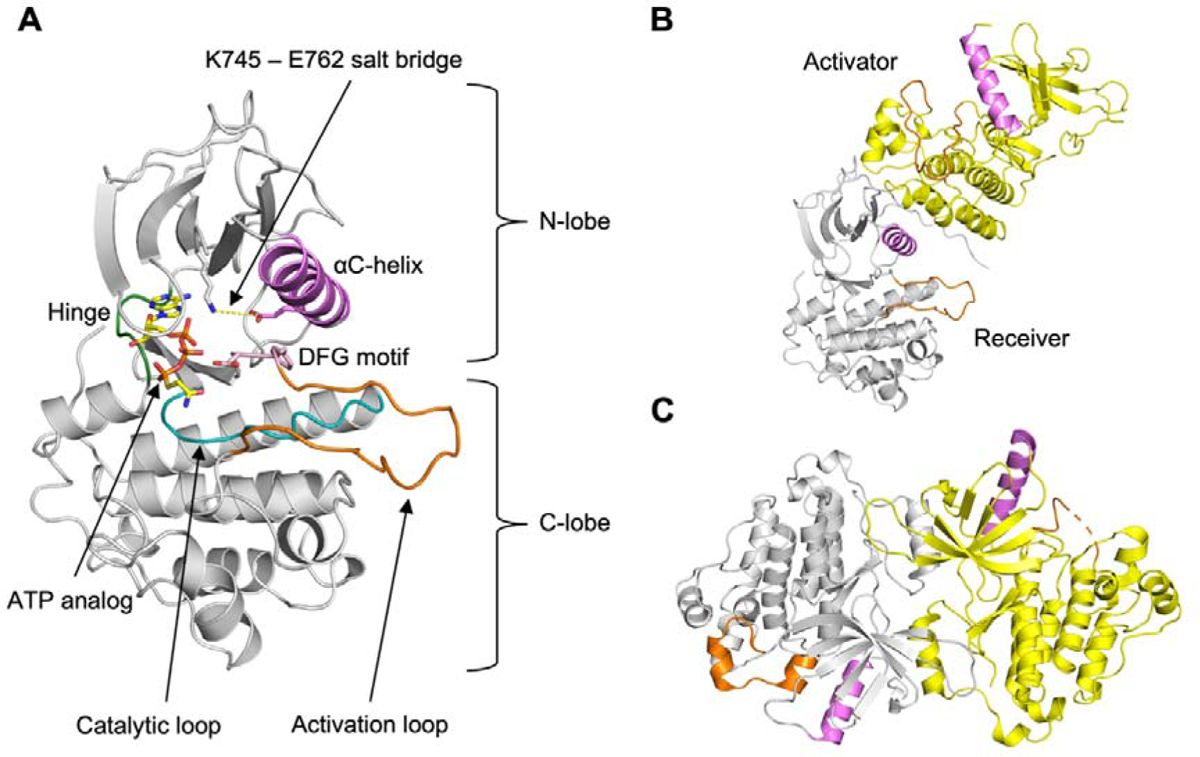
Structure of EGFR in monomeric and dimeric form. (A) X-ray structure of the EGFR kinase domain in active conformation (PDB: 2GS6), with key structural features highlighted. (B) The kinase domain can form an asymmetric dimer, where both protomers adopt an active-like conformation. The receiver and activator chains are colored gray and yellow, respectively. (C) EGFR can also form a symmetric dimer, in which both chains adopt the Src-like inactive conformation (PDB: 6V66). In this arrangement, the activation loop is collapsed. The chains are colored gray and yellow, respectively. For better comparison, the αC-helices (violet) and activation loops (orange) are highlighted in both (B) and (C).

EGFR kinase domains form distinct intracellular dimer configurations that regulate activity. In the asymmetric dimer (Figure 1b), both chains adopt the active conformation [10]. One chain, the activator, presses against the other, the receiver, locking the receiver’s αC-helix in the active orientation. This dimerization mechanism is a crucial pre-requisite for the activation of wild-type EGFR under physiological conditions. Conversely, the symmetric dimer (Figure 1c) consists of two chains in the Src-like inactive conformation, stabilized by interactions between their C-terminal tails, and is believed to play a role in EGFR autoinhibition. Apart from the allosteric effects of dimerization on EGFR activity, mutations in the kinase domain can also modulate EGFR conformation and activity.

More than 500 different EGFR missense mutations have been recorded in the Catalogue of Somatic Mutations in Cancer (COSMIC) database. The exon 21 mutation L858R, along with exon 19 deletion mutations, is among the most frequent oncogenic driver mutations in EGFR-positive NSCLC [11]. Its role in causing overactivation and its response to targeted therapy have been extensively discussed in the literature. L858R is located at the beginning of the activation loop, adjacent to the DFG motif, and has been found to both stabilize the active conformation and disrupt the inactive conformation [6, 12–14]. Patients treated with TKIs for L858R-positive NSCLC frequently acquire a secondary resistance mutation, T790M, leading to the double mutation T790M/L858R. Although historically termed the ‘gatekeeper’ mutation, T790M/L858R confers drug resistance not through steric hindrance but likely by elevating ATP affinity [15].

Beyond these well-characterized mutations, hundreds of rare and ultra-rare EGFR mutations exist, many of which are not yet biochemically characterized, and for most, the structural mechanisms remain unknown. The rare exon 20 mutation S768I, accounting for approximately 0.6 – 1% of EGFR mutations in NSCLC, has been well established as oncogenic in both experimental and clinical studies [16–18]. Located at the C-terminal end of the αC-helix and facing away from the binding site, S768I has been proposed by Shan et al. (identified as S744I in their study) to stabilize the active conformation through an allosteric mechanism [6]. In their simulations, they detected prolonged K745 – E762 salt bridge formation and increased αC-helix stability compared to the wild-type, and attributed these changes to enhanced hydrophobic packing between the mutated isoleucine residue in the αC-helix and the activation loop. However, their study did not quantify this increased hydrophobic packing. In contrast, an ultra-rare variant, S768N, located at the same position, has been experimentally suggested as a neutral mutation similar to the wild-type [19, 20]. The structural mechanism underlying S768N’s non-activating nature, however, remains unknown.

Another αC-helix mutation, the exon 19 mutation D761N, has been identified in multiple tumor samples. It has been associated with increased EGFR activity and resistance to TKIs [20–22]. Like S768N, its structural mechanism remains unknown, though its location distal from the ATP-binding site suggests it alters EGFR activity through an allosteric mechanism.

Molecular Dynamics (MD) simulations are widely used to investigate the structural mechanisms of EGFR mutations by comparing mutant dynamics to wild-type EGFR. Previous EGFR simulation studies have employed diverse system configurations and protocols, varying in ATP inclusion, dimerization states, kinase domain conformation, simulation time, and force field parameters [1, 6–9, 12, 14, 23–30]. To the best of our knowledge, no benchmarking study has been reported that establishes optimal simulation parameters specifically tailored for EGFR and its mutants. As a result, existing MD studies on EGFR may lack comparability unless the simulation protocols are clearly communicated and standardized. A further challenge is that many previous MD studies rely on long, microsecond-scale simulations, which typically require weeks to complete and involve extensive analysis. Consequently, long simulation-guided EGFR studies are often constrained by computational resources, limiting their applicability to a broader range of mutants.

To address these challenges, we developed a standardized protocol for rapidly assessing EGFR mutants using structural modeling and multiple short MD simulations. We compared the popular CHARMM and AMBER force fields through simulations of wild-type, L858R, and the double mutant T790M/L858R as well-characterized reference systems. We then applied this protocol to the rare mutations S768I, S768N, and D761N. Our simulation results were consistent with the known pathogenic behavior of these variants and revealed distinct mechanisms by which EGFR activation is affected. Using our protocol, we could reproduce the previously reported increases in K745 – E762 salt bridge occupancy and αC-helix stability for the oncogenic S768I mutant. We also expanded upon previous findings by quantifying the proposed increase in hydrophobic interactions formed by S768I. In contrast, the hypothetically neutral mutation S768N increased local disorder by altering its hydrogen bonding pattern, forming less stable hydrogen bonds than wild-type, and exhibited reduced K745 – E762 salt bridge formation compared to S768I. Meanwhile, the oncogenic D761N mutation reshaped the hydrogen bonding network, introducing a novel and stable interaction pattern between the αC-helix and the activation loop. Altogether, our study demonstrates that this MD protocol serves as a powerful tool to predict the oncogenic potential of EGFR mutations while elucidating their molecular modes of action. Importantly, our protocol does not rely on microsecond-scale simulations, making it suitable for efficiently screening EGFR mutations (e.g. < 2 days).

## 2. Materials and Methods

### 2.1 Structural modeling

The initial structures for the simulation of EGFR kinase domain dimers were generated using the homology modeling software MODELLER (version 10.4) [31], based on a template structure in the active conformation (PDB:2GS6) for the asymmetric dimer and a template structure in the inactive conformation (PDB:6V66) for the symmetric dimer. ATP + Mg^2+^ were placed in the ATP-binding pockets of the template structures to obtain ATP-bound templates. The canonical EGFR UniProt sequence (AC:P00533), from residue 697 to 1022, was selected as the target sequence for the wild-type. Mutations were introduced into the wild-type sequence at the appropriate location to generate the target sequences for the mutants. For each target sequence, 2000 models were generated for both the asymmetric and symmetric ATP-dimer complexes, and the best model from each set was selected based on the DOPE score [32]. The quality of the generated models was evaluated using MolProbity [33]. Apo structures were obtained by removing ATP + Mg^2+^ from the models.

### 2.2 Molecular Dynamics (MD) simulations

All simulations were performed with GROMACS (versions 2021.5 and 2022.1) [34], using the CHARMM36m force field [35] for CHARMM-based simulations and the AMBER ff19SB force field [36] for AMBER-based simulations. The input files for the simulations were generated with the CHARMM-GUI web interface (https://www.charmm-gui.org) [37–40]. The N-terminus (-NH_2_) and C-terminus (-COOH) of EGFR were capped by acetylation and amidation, respectively. Histidine residues were treated as δ-nitrogen protonated, except for histidine 835, which was treated as ε-nitrogen protonated. In the holo simulations, the ATP molecule was assigned a 4-fold negative charge. Each EGFR dimer was solvated in a rectangular box with an edge distance of 10.0 Å from the solute, and the total charge of each system was neutralized by the addition of Na^+^ and Cl^-^ ions to a final concentration of 0.15 M. Systems were energy-minimized using the steepest descent algorithm and then equilibrated for 125 ps under NVT conditions at 303.15 K. Production runs under NPT conditions were subsequently carried out in 100 ns chunks (‘replicas’) with randomly initialized velocities. Each system was simulated with five independent replicas, amounting to 500 ns per system (see also Tables S1 and S2 in the Supplementary Material).

### 2.3 Data analysis and visualization

Averages are presented as mean ± standard deviation. Simulation trajectories were analyzed using built-in tools from GROMACS and the Python library MDTraj [41]. The K745 – E762 salt bridge was considered intact when the distance between the NZ atom of lysine and either of the carboxylate oxygens (OE1 or OE2) of glutamate was within 4.0 Å. Activation loop disruption was defined as occurring when the one-turn helix pitch exceeded a threshold of 7.0 Å. The helix pitch was measured as the distance between the alpha carbon (Cα) atoms of residues 858 and 862. Protein structures were visualized using the PyMOL Molecular Graphics System, Version 3.0.2 Schrödinger, LLC.

## 3. Results

### 3.1 Benchmarking of the Simulation Protocol

To establish an effective MD protocol for studying dynamic differences between wild-type and mutant EGFR, we compared two widely used force fields, CHARMM and AMBER. We simulated both the active-like and Src-like inactive conformations (hereafter referred to as ‘active’ and ‘inactive’ conformations, respectively). As discussed earlier, the active conformation is associated with the asymmetric dimer, whereas the inactive conformation can form a symmetric dimer. Accordingly, we simulated each conformation in its respective dimeric state. To assess the effect of ATP on EGFR dynamics, we simulated each dimer in both apo (no ligand present) and holo (ATP bound in the receiver monomer) conditions. Since the symmetric dimer lacks a defined receiver chain, ATP was placed arbitrarily in the first chain (chain ‘A’) as defined in the crystal structure assignment.

For benchmarking, we simulated wild-type (WT) EGFR, the oncogenic driver mutant L858R, and the common resistance double mutant T790M/L858R (TMLR). Each combination of force field, dimer type, apo/holo state, and mutation/wild-type status constituted a distinct simulation system. Each system was simulated for 500 ns. With 24 systems in total, our benchmarking study comprised 12 µs of simulation time (Table S1).

### 3.2 CHARMM vs. AMBER: Simulating the Asymmetric Dimer

A well-established model to explain EGFR overactivation involves the stabilization of its active conformation [6, 12]. This active conformation is characterized by the formation of the catalytically important K745 – E762 salt bridge (Figure 2a). To assess the stability of the active state, we measured the occupancy of this salt bridge in our simulations of the asymmetric dimer (Figure 2b).

**Figure 2.**
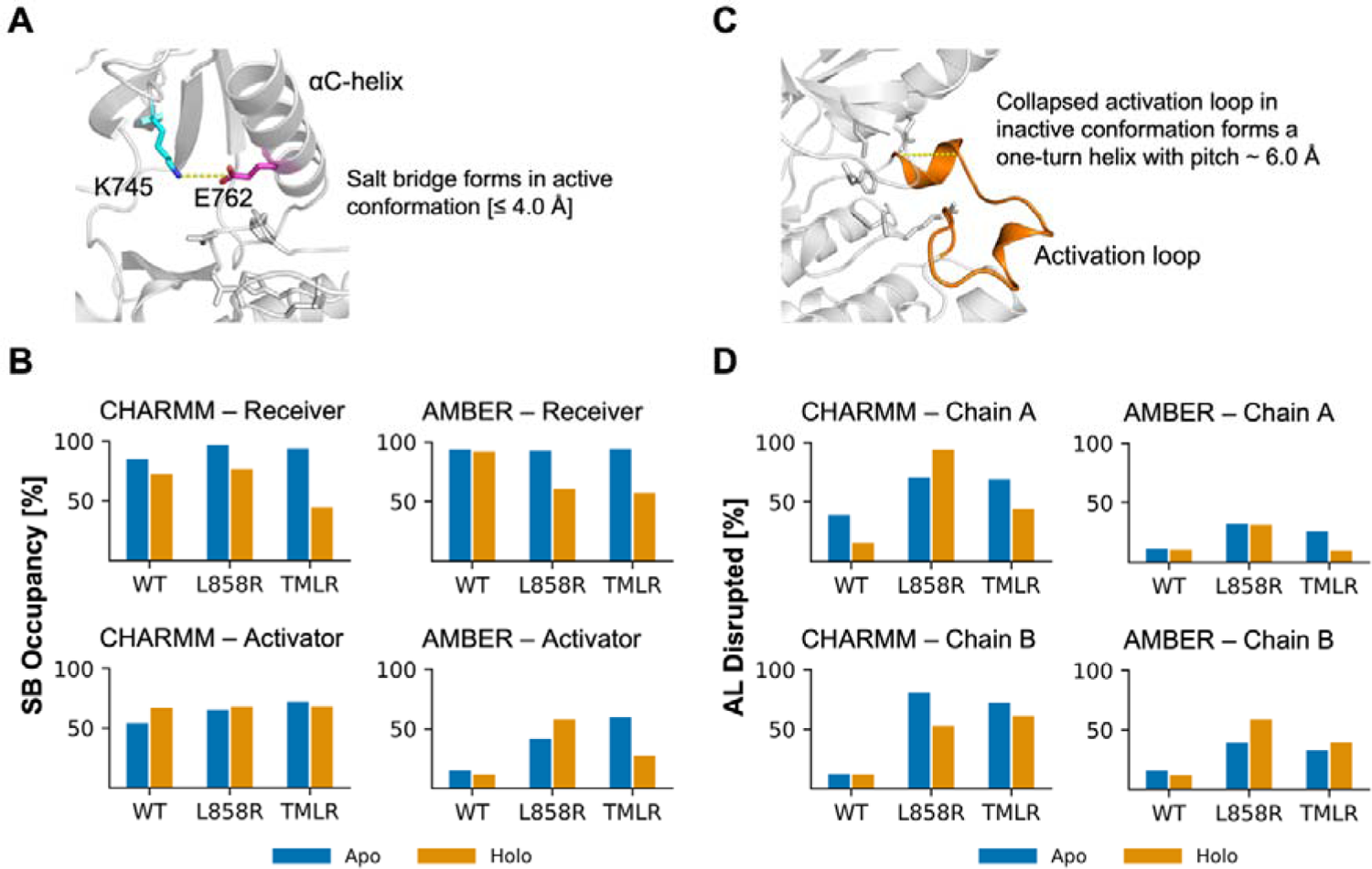
Benchmarking the CHARMM and AMBER force fields for reproducing known EGFR dynamics. (A) In the active conformation, a salt bridge forms between residues K745 and E762. (B) Salt bridge occupancy using a cutoff of 4.0 Å in simulations of the asymmetric dimer. Values are shown for different force fields, chains, and the presence or absence of ATP. Note that data are presented only for the asymmetric dimer, as the K745 – E762 salt bridge did not form in simulations of the symmetric dimer. (C) In the Src-like inactive conformation, the activation loop is collapsed, forming a short, one-turn helix with a pitch of approximately 6.0 Å. (D) Disruption of the inactive activation loop conformation. The loop was considered disrupted if the helix pitch exceeded 7.0 Å. The percentage of simulation time during which the loop was disrupted is shown for different force fields, chains, and the presence or absence of ATP. Note that data are presented only for the symmetric dimer, as the one-turn helix was not observed in simulations of the asymmetric dimer. Abbreviations: SB = salt bridge, AL = activation loop.

In CHARMM simulations of the apo state, salt bridge occupancy was increased for both mutants relative to WT in both chains of the asymmetric dimer. In the receiver chain, this effect was particularly pronounced for L858R, which showed an increase of more than 10 percentage points compared to WT (WT: 85.4%, L858R: 97.1%, TMLR: 93.8%). In the apo activator chain, TMLR exhibited a strong increase in salt bridge occupancy of nearly 20 percentage points, while L858R also showed a substantial increase of over 10 percentage points relative to WT (WT: 53.9%, L858R: 65.1%, TMLR: 72.0%).

In contrast, AMBER apo simulations showed relatively consistent salt bridge occupancy in the receiver chain across variants (WT: 93.8%, L858R: 92.6%, TMLR: 94.1%). In the activator chain, however, occupancies were generally lower than those observed in CHARMM and exhibited greater variation across mutants. Notably, TMLR showed the highest occupancy in the apo activator chain among all variants, with an occupancy more than 40 percentage points higher than WT (WT: 15.0%, L858R: 41.6%, TMLR: 59.8%).

In CHARMM holo simulations of the asymmetric dimer, salt bridge occupancy in the receiver chain was slightly higher for L858R than in WT (WT: 72.6%, L858R: 76.7%), but was markedly reduced for TMLR (44.4%). In the activator chain, occupancies were similar across all three variants (WT: 66.8%, L858R: 67.7%, TMLR 67.9%).

In AMBER holo simulations, both L858R and TMLR exhibited lower salt bridge occupancy in the receiver chain compared to WT, with reductions of more than 30 percentage points (WT: 92.0%, L858R: 60.4%, TMLR: 57.0%). In contrast, occupancy in the activator chain increased substantially for both mutants relative to WT, with a particularly pronounced increase for L858R (WT: 11.4%, L858R: 57.6%, TMLR: 27.6%).

Several key trends emerged from these simulations. Within each CHARMM simulation condition, L858R consistently exhibited higher salt bridge occupancy than WT, suggesting that it stabilizes the active EGFR conformation. In AMBER simulations, this increase was observed only in the activator chain but not in the receiver chain. Additionally, the difference in salt bridge occupancy between WT and mutant activator chains was larger in AMBER simulations than in CHARMM, primarily due to the near absence of salt bridge formation in the WT activator chain in AMBER. Across both force fields, salt bridge occupancy in the receiver chain decreased upon ATP binding compared to apo conditions, likely due to the displacement of the E762 carboxylate group by ATP’s phosphate moiety, causing K745 to interact with ATP rather than E762. Consequently, salt bridge occupancy in the holo receiver chain may not reliably indicate stabilization of the active conformation.

### 3.3 CHARMM vs. AMBER: Simulating the Symmetric Dimer

An alternative mechanism proposed for EGFR overactivation involves the destabilization of the inactive, symmetric dimer [12, 13]. In this model, mutations that reduce compatibility with the inactive conformation shift the conformational equilibrium toward the active state, potentially resulting in gain of function effects.

The Src-like inactive conformation is characterized by a short, one-turn helix at the beginning of the activation loop, immediately following the DFG motif (Figure 2c). Consistent with previous studies, our simulations of the symmetric dimer showed that mutants disrupted this helix more frequently than WT. To quantify this effect, we measured helix disruption in the activation loop as a proxy for destabilization of the inactive state.

In CHARMM apo simulations of the symmetric dimer, activation loop disruption was observed in 38.7% and 12.1% of the simulation time in chains A and B for WT, respectively. In contrast, L858R exhibited a substantial increase in disruption, occurring in 70.6% of the simulation time in chain A (an increase of over 30 percentage points relative to WT) and 80.6% of the time in chain B (an increase of nearly 70 percentage points relative to WT; Figure 2d). TMLR showed similar trends, with activation loop disruption occurring 69.0% and 72.3% of the time in chains A and B, respectively, under CHARMM apo conditions.

AMBER apo simulations of the symmetric dimer also demonstrated increased activation loop disruption in mutants compared to WT, though the effect was less pronounced than in CHARMM. For WT, activation loop disruption occurred in 11.2% and 15.8% of the simulation time in apo chains A and B, respectively. In contrast, L858R showed increases of more than 20 percentage points relative to WT, with disruption observed 32.1% of the time in chain A and 39.2% in chain B. TMLR exhibited a more moderate increase, with disruption observed 25.8% and 32.7% of the time in chains A and B, respectively.

In CHARMM holo simulations of the symmetric dimer, activation loop disruption increased sharply for L858R and TMLR compared to WT. While WT exhibited disruption in only 15.3% and 11.8% of the simulation time in chains A and B, respectively, L858R showed a dramatic rise, with disruption occurring 93.8% and 52.5% of the time. TMLR also displayed increased disruption, at 44.3% and 61.1% in chains A and B, respectively.

In AMBER holo simulations, activation loop disruption in WT was observed 10.5% and 11.6% of the time in chains A and B, respectively. While L858R exhibited increased disruption in both chains (31.6% and 58.4% of the time), TMLR showed no increase in chain A (9.9% of the time) but a substantial increase in chain B (39.4% of the time).

Overall, both CHARMM and AMBER simulations demonstrated that L858R and TMLR disrupt the inactive-like activation loop conformation more than WT. This effect was more pronounced in CHARMM, where the absolute increase in disruption was greater. While full transition to an extended activation loop conformation was not observed, the disruption of the activation loop helix suggests that L858R and TMLR destabilize the inactive state.

### 3.4 Evaluation of the benchmarking study

We then evaluated which protocol to use for simulating additional variants. While both force fields produced results that generally aligned with existing findings for WT and mutants, we opted to continue using only the CHARMM force field for two main reasons. First, CHARMM more consistently captured the expected increase in K745 – E762 salt bridge stability of L858R relative to WT in asymmetric dimer simulations (Figure 2b). While AMBER showed a more pronounced increase in salt bridge occupancy in the activator chain for L858R relative to WT compared to CHARMM, it showed no increase in occupancy in the receiver chain. Second, CHARMM better captured the destabilization of the inactive conformation by the cancer mutants (Figure 2d). In CHARMM simulations, L858R and TMLR caused a more pronounced disruption of the inactive activation loop conformation relative to WT, compared to AMBER simulations. We therefore concluded that using the CHARMM force field alone is sufficient to capture mutational overactivation.

We then applied this MD protocol to study the rare mutations S768I, S768N, and D761N. Each mutant was simulated in asymmetric and symmetric dimeric forms under apo and holo conditions with the CHARMM force field for 500 ns per system, totaling 2 µs per mutant (Table S2).

### 3.5 S768I exhibits increased hydrophobic packing between **α**C-helix and activation loop

The oncogenic mutation S768I has been reported to stabilize the αC-helix and enhance K745 – E762 salt bridge formation compared to the wild-type. This stabilization has been attributed to increased hydrophobic packing introduced by the mutation. To assess whether our simulations reproduced these findings, we analyzed salt bridge formation, secondary structure stability, and the number of hydrophobic contacts formed by residue 768. Additionally, we investigated whether S768I disrupts the inactive conformation.

As expected, apo simulations of the asymmetric dimer confirmed that the K745 – E762 salt bridge is more stable in S768I than in WT (Figure 3a): the salt bridge forms 96.6% of the time in the receiver chain of S768I (WT: 85.4%, for comparison: L858R: 97.1%) and 78.4% of the time in the activator chain, which is more than 20 percentage points higher than WT (WT: 53.9%, L858R: 65.1%). In the holo simulations of the asymmetric dimer, S768I exhibited decreased salt bridge formation (51.9%) relative to WT in the receiver chain (WT: 72.6%, L858R: 76.7%), but increased formation (78.3%) in the activator chain (WT: 66.8%, L858R: 67.7%). To determine whether the mutated residue directly interacts with the salt bridge residues, we measured its proximity to key catalytic sites. In all simulations, the sidechain heavy atoms of both WT and mutant residue 768 remained beyond 7 Å from the side chains of K745 and E762, 10 Å from the sidechain of the catalytic aspartate D855 of the DFG motif, and 13 Å from any ATP heavy atom. Therefore, residue S768 does not directly interact with ATP or catalytically relevant residues, suggesting an allosteric mechanism for the increase in salt bridge formation.

**Figure 3.**
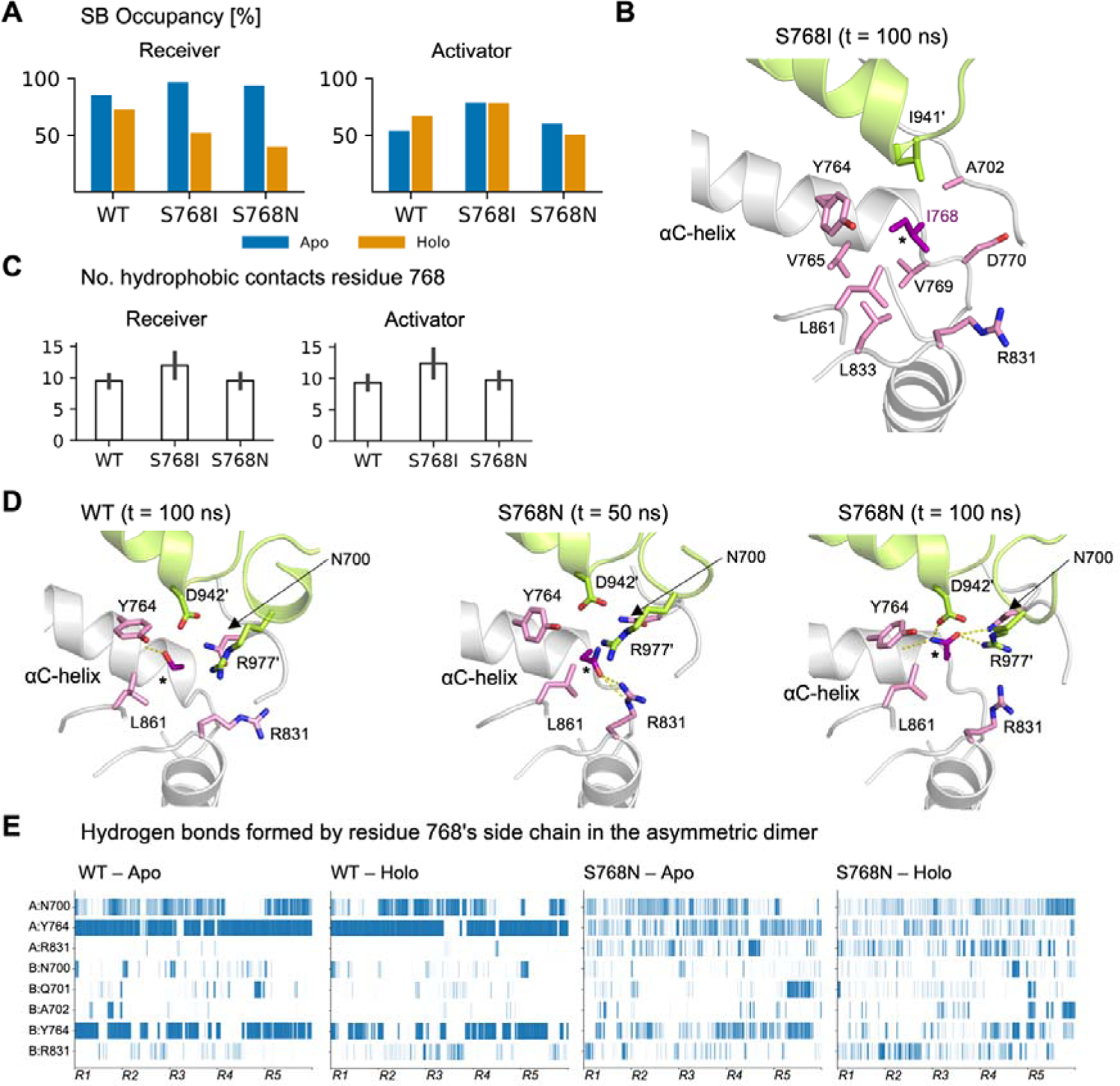
Allosteric regulation in the mutant pair S768I and S768N. (A) Comparison of salt bridge occupancy in the asymmetric dimer among WT, S768I, and S768N, analogous to Figure 2b. (B) Snapshot of the S768I mutant in the asymmetric dimer conformation after 100 ns of simulation time. The mutant residue I768 and surrounding residues form a hydrophobic network. The receiver and activator chains are highlighted in light gray and green, respectively. I768 is highlighted in purple and labeled with an asterisk. Other residues of the receiver chain are shown in pink stick representation, while residues of the activator chain are depicted in green stick representation. (C) Average number of hydrophobic contacts between residue S/I/N768 and its neighboring residues in the asymmetric dimer, calculated across both Apo and Holo systems. Error bars show standard deviation. (D) Snapshots of WT and the S768N mutant at different simulation times. Hydrogen bonds involving the sidechain of residue 768 are shown as yellow dashed lines. The receiver and activator chains are highlighted in light gray and green, respectively. Residue 768 is highlighted in purple and labeled with an asterisk. Other residues of the receiver chain are shown in pink stick representation, while residues of the activator chain are depicted in green stick representation. (E) Hydrogen bond existence map illustrating interactions between the sidechain of residue 768 and its surrounding environment over time. Each residue is labeled by its chain: A = receiver chain, B = activator chain. Only the most frequently observed hydrogen bonds are depicted. Data from all replicas are combined for each system. Abbreviations: SB = salt bridge, Rep = Replica.

To investigate the reported stabilization of the active conformation of the αC-helix, we assessed its secondary structure stability using the DSSP method. Consistent with previous findings, S768I exhibited greater αC-helix stability in the asymmetric dimer compared to WT, with a higher average number of residues assigned as ‘helical’ in both apo and holo simulations. In the apo state, this increase was observed in both the receiver (S768I: 15.4 ± 1.1, WT: 15.0 ± 1.2) and activator (S768I: 15.4 ± 1.1, WT: 15.1 ± 1.4) chains. A similar trend was seen in holo simulations, with S768I maintaining a higher number of helical residues in both the receiver (S768I: 15.7 ± 0.8, WT: 14.6 ± 1.4) and activator (S768I: 14.9 ± 1.5, WT: 14.7 ± 1.5) chains.

To quantify the reported increase in hydrophobic packing, we measured hydrophobic contacts between residue 768 and its neighboring residues (Figure 3b,c). Supporting previous claims, S768I exhibited approximately 30% more hydrophobic interactions than WT. This increase was observed in all conditions: apo receiver chain (WT: 9.4 ± 0.9, S768I: 11.8 ± 2.0), apo activator chain (WT: 9.4 ± 1.1, S768I: 12.3 ± 2.3), holo receiver chain (WT: 9.4 ± 1.0, S768I: 12.1 ± 2.0), and holo activator chain (WT: 9.2 ± 1.0, S768I: 12.5 ± 2.3).

We also analyzed the disruption of the short one-turn helix in the collapsed activation loop in simulations of the inactive, symmetric dimer. Compared to WT, S768I exhibited reduced activation loop disruption in apo chain A (WT: 38.7%, S768I: 27.8%) but increased disruption in apo chain B (WT: 12.1%, S768I: 38.4%), holo chain A (WT: 15.3%, S768I: 16.5%), and holo chain B (WT: 11.8%, S768I: 27.6%).

However, even in systems where S768I increased disruption of the inactive state, the effect was less pronounced than in L858R or TMLR (Figure 2d and Table S3), suggesting that S768I does not strongly destabilize the inactive state.

In summary, our simulations of the asymmetric dimer showed that S768I increases salt bridge formation under apo conditions, enhances αC-helix stability, and strengthens hydrophobic packing. Notably, the increase in salt bridge formation was comparable to or greater than that observed for the strongly oncogenic L858R under all conditions, except in the receiver chain in holo simulations, where – as mentioned in section 3.2 – salt bridge stability may not be an indicative measure of EGFR overactivation. The increase in hydrophobic interactions between the mutated residue 768 and hydrophobic residues from both the αC-helix and activation loop (Figures 3b,c) likely contributes to the observed stabilization of the αC-helix. Given that E762, one of the two residues involved in the catalytic salt bridge, is part of the αC-helix, this stabilization may play a role in reinforcing the salt bridge. Since S768I did not induce as strong a disruption of the inactive state as L858R and TMLR, we hypothesize that its oncogenic effects primarily stem from stabilizing the active state rather than destabilizing the inactive conformation.

### 3.6 S768N increases local disorder through formation of a dynamic hydrogen bonding network

To investigate the structural basis for the presumed non-activating behavior of the S768N mutant, we performed the same analyses as for the oncogenic S768I mutant. As expected, S768N exhibited reduced salt bridge formation and weaker hydrophobic packing compared to S768I (Figures 3a,c) in the asymmetric dimer. Additionally, αC-helix stability in S768N was either lower or comparable to WT. In the apo state, this reduction was observed in both the receiver (S768N: 14.2 ± 1.2, WT: 15.0 ± 1.2) and activator (S768N: 14.5 ± 1.5, WT: 15.1 ± 1.4) chains. A similar trend was seen in holo simulations, where S768N maintained comparable or reduced numbers of helical residues in both the receiver (S768N: 14.7 ± 1.5, WT: 14.6 ± 1.4) and activator (S768N: 14.3 ± 2.0, WT: 14.7 ± 1.5) chains.

We also analyzed the disruption of the collapsed activation loop in the symmetric dimer. Compared to WT and S768I, S768N exhibited increased activation loop disruption. However, this effect was less pronounced than in L858R or TMLR (Table S3). These findings suggest that S768N is unlikely to confer a gain of function, supporting its classification as a neutral mutation. To further explore why the structural changes caused by S768N were likely insufficient to drive overactivation, we investigated their impacts on the local hydrogen bonding network.

We hypothesized that the asparagine substitution in S768N might alter hydrogen bonding patterns, thereby affecting the stability of the active conformation. To test this, we analyzed hydrogen bonding interactions involving the serine side chain in WT and the asparagine side chain in S768N (Figure 3d). Interestingly, while S768N is theoretically capable of forming more hydrogen bonds – at times forming up to five simultaneously in our simulations – the average number of hydrogen bonds observed across simulation snapshots was equal to or lower than in WT. In apo simulations of the asymmetric dimer, the WT serine side chain formed an average of 1.4 ± 0.6 and 0.9 ± 0.7 hydrogen bonds in the receiver chain and the activator chains, respectively. In contrast, the S768N side chain formed 0.9 ± 0.7 and 0.8 ± 0.9 hydrogen bonds in the receiver and activator chains, respectively. Similar trends were observed in holo simulations, where WT formed 1.3 ± 0.5 and 0.8 ± 0.6 hydrogen bonds, while S768N formed 0.8 ± 0.7 and 0.9 ± 0.9 hydrogen bonds in the receiver and activator chains, respectively.

Next, we examined the hydrogen bonding partners of residue 768 in our asymmetric dimer simulations (Figure 3e). In WT, hydrogen bonds were primarily formed with the backbone carbonyl oxygen of the αC-helix residue Y764 and, less frequently, with the side chain of N700 in the N-terminal tail. In contrast, the side chain of S768N exhibited greater flexibility, frequently switching hydrogen bonding partners between N700, Y764, and R831 (located in the αE-helix of the C-lobe), with occasional interactions with Q701 and A702. We propose that this increased flexibility, which resulted in an overall reduction in hydrogen bond stability due to numerous short-lived interactions, contributes to greater disorder in the αC-helix and may explain the presumed non-activating behavior of S768N. Interestingly, in the inactive, symmetric dimer, S768N formed slightly more hydrogen bonds than WT (Table S4), potentially suggesting a modest stabilizing effect on the inactive state.

### 3.7 D761N forms a new hydrogen bonding pattern between the **α**C-helix and activation loop

We also simulated the oncogenic exon 19 mutation D761N, located near the center of the αC-helix. When measuring salt bridge occupancy in the apo asymmetric dimer, we found that D761N exhibited the highest occupancy among all simulated mutants in the receiver chain, increasing by 12 percentage points compared to WT (from 85.4% to 97.5%, Figure 4a). In contrast, salt bridge occupancy decreased in the activator chain (WT: 53.9%, D761N: 49.3%). This suggests that D761N primarily stabilizes the active conformation in the receiver chain but not in the activator chain. Additionally, D761N did not increase activation loop disruption relative to WT in the inactive, symmetric dimer (Table S3). Therefore, similar to S768I, our simulations suggest that D761N primarily stabilizes the active state rather than destabilizing the inactive state.

**Figure 4.**
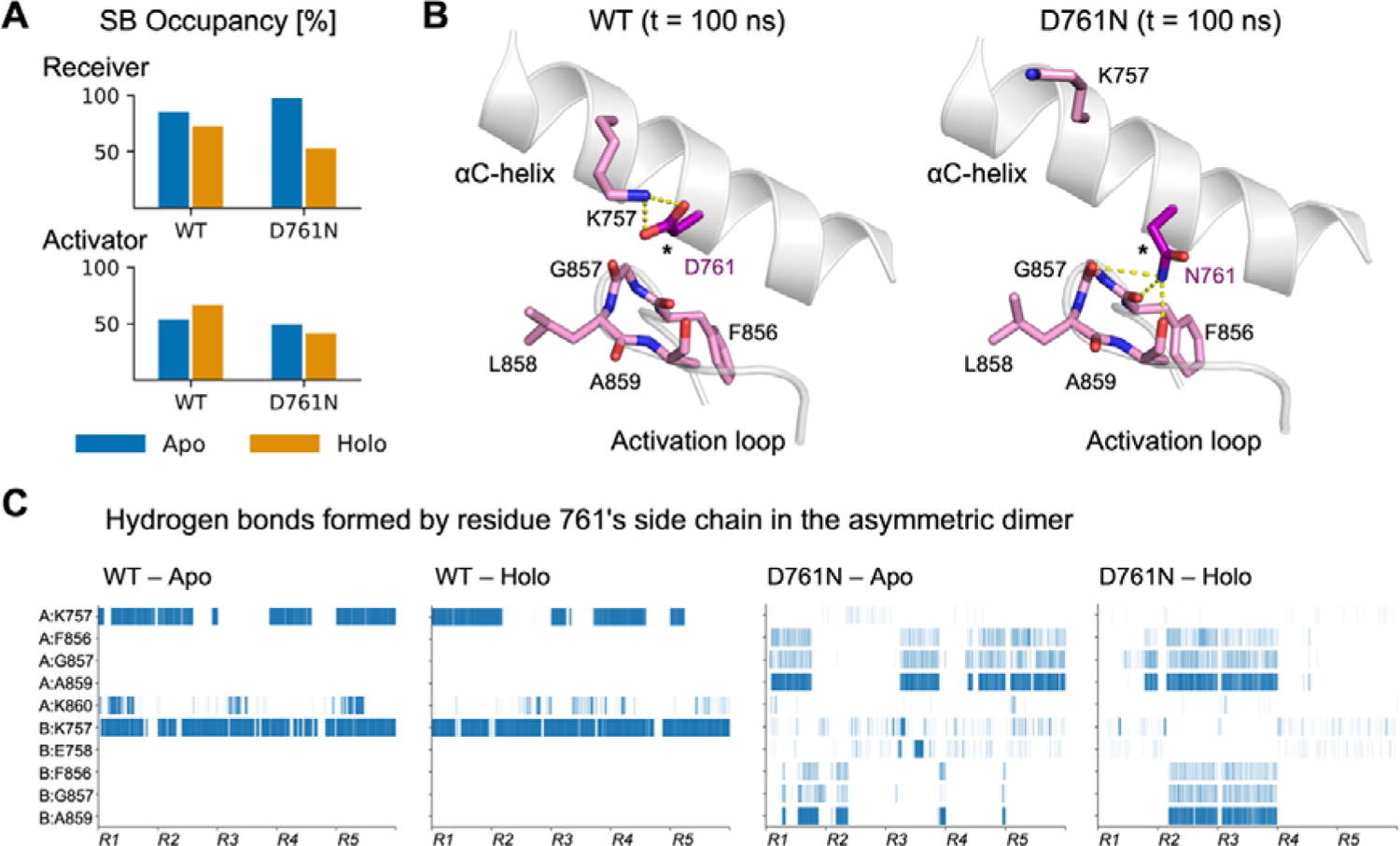
The D761N mutation forms a new hydrogen bonding pattern between the αC-helix and the activation loop. (A) Comparison of salt bridge occupancy in the asymmetric dimer among WT and D761N, analogous to Figure 2b. (B) Snapshots of WT and the D761N mutant after 100 ns of simulation time each. Hydrogen bonds involving the sidechain of residue 761 are shown as yellow dashed lines. The snapshots display the receiver chain, with residue 761 highlighted in purple and marked with an asterisk. For residues 856-859, the backbone atoms are also shown. (C) Hydrogen bond existence map illustrating interactions between the sidechain of residue 761 and its surrounding environment over time. Each residue is labeled by its chain: A = receiver chain, B = activator chain. Only the most frequently observed hydrogen bonds are depicted. Data from all replicas are combined for each system. Abbreviations: SB = salt bridge, R = Replica.

To further explore the structural effects of D761N, we examined the local hydrogen bonding network around the mutant site (Figure 4b). Interestingly, D761N formed a stable alternative hydrogen bonding network distinct from that of WT. In WT, the aspartate side chain of D761 primarily formed a salt bridge with the more N-terminal lysine K757. In contrast, the mutated asparagine side chain in the D761N mutant frequently engaged in hydrogen bonds with the backbones of residues F856, G857, and A859 (Figure 4c). Notably, F856 and G857 belong to the DFG motif, and all three residues are part of the activation loop. In the apo state, these hydrogens bonds formed more frequently in the receiver chain than in the activator chain, aligning with the high salt bridge stability observed for the apo receiver chain.

Conversely, under holo conditions, the hydrogen bond formation was more balanced between the receiver and activator chains, which also corresponded to the similar salt bridge occupancy between the holo receiver and activator chains (Figure 4a). These findings suggest that D761N stabilizes a novel hydrogen bonding pattern between the αC-helix and the activation loop, potentially promoting the active conformation allosterically and contributing to oncogenicity.

## 4. Discussion

### 4.1 A novel protocol to study EGFR overactivation

Long-timescale unbiased MD simulations have been widely employed to study EGFR conformational transitions, with some studies extending into the hundreds of microseconds of simulation time [6, 7]. Others have used enhanced sampling methods to explore a broader conformational space [8, 12]. Here, we presented a protocol that utilizes sets of 500 ns unbiased simulations to compare the dynamics of wild-type EGFR and its mutants. This timescale is sufficient to observe disruptions in the active EGFR state [6]. Although the Src-like inactive state is relatively stable – having been simulated for up to 100 µs in previous studies – our analysis of activation loop stability in the inactive, symmetric dimer captured local disruptions well within our 500 ns simulation time frame. To ensure reproducibility in system preparation and simulation setup, we used the well-established CHARMM-GUI web interface (see Methods). This approach standardizes the simulation workflow, particularly for extending studies to additional EGFR variants. Furthermore, our analysis steps are easily repeatable and can serve as a blueprint for studying other EGFR mutations or even other kinase family members.

By simulating both the active, asymmetric dimer and the inactive, symmetric dimer, our protocol enables identification of whether a mutation primarily affects the active state, the inactive state, or both. This distinction has practical implications for drug design, as different classes of tyrosine kinase inhibitors (TKIs) target specific EGFR conformations [42]. Notably, Type I TKIs bind to the active state, while Type I.5 TKIs preferentially target the Src-like inactive state. Understanding the mechanism of overactivation, as revealed by our protocol, could help guide the selection of therapeutic strategies by identifying which conformational state to target.

### 4.2 Applicability to further variants

We aimed to establish criteria to determine whether a mutation is likely oncogenic based on our protocol. Our results suggest that CHARMM apo simulations alone are sufficient to distinguish between oncogenic and neutral mutants. Across these simulations, we observed that all oncogenic mutations examined in this study (L858R, TMLR, S768I, and D761N) increased K745 – E762 salt bridge occupancy by at least 10 percentage points in at least one chain of the apo asymmetric dimer, relative to WT, which exhibited occupancies of 85.4% and 53.9% in the receiver and activator chains, respectively. Notably, L858R and S768I met this threshold in both chains. In addition, TMLR and L858R were the only oncogenic mutants that consistently exhibited high disruption of the inactive state. Under apo conditions, both mutations disrupted the inactive conformation in more than 60% of the simulation time for both chains – a threshold that no other mutant exceeded in apo simulations. Based on these observations, a mutation is likely oncogenic if it meets at least one of the following three criteria in apo simulations: (i) salt bridge occupancy of ∼95% or higher in the receiver chain of the asymmetric dimer, (ii) salt bridge occupancy of ∼64% or higher in the activator chain of the asymmetric dimer, (iii) disruption of the inactive state in at least 60% of the simulation time for both chains in the symmetric dimer. We suspect that the likelihood of oncogenic hyperactivation increases when multiple criteria are met. Additionally, if ambiguity remains, holo simulations may provide further insights.

## 5. Conclusion

The EGFR kinase domain exhibits a vast mutational landscape, with nearly 1,000 recorded somatic mutations cataloged in COSMIC, including over 500 missense mutations. Despite this diversity, only a small fraction of these mutants has been biochemically characterized, and even fewer have been studied at the structural level. Our MD protocol successfully captured the differences between wild-type EGFR and the oncogenic and neutral mutations studied here. For the frequent oncogenic mutations L858R and T790M/L858R, our simulations demonstrated both stabilization of the active state and disruption of the inactive state, consistent with their known oncogenic mechanisms. In contrast, the rare oncogenic mutations S768I and D761N stabilized the active conformation, with our simulations suggesting distinct allosteric mechanisms as the underlying cause – either through increased hydrophobic packing around the αC-helix (S768I) or the formation of a new hydrogen bonding network between the αC-helix and activation loop (D761N). Meanwhile, the neutral mutation S768N failed to stabilize the active conformation, likely due to an unstable and overly flexible hydrogen bonding network introduced by the mutation.

Importantly, our study demonstrates that even relatively short 500 ns simulations – which can typically be completed within two days on modern computational hardware – are sufficient to capture key mutational effects on EGFR overactivation. These results highlight the potential of MD simulations for rapidly screening EGFR mutations and gaining insights into their structural and functional consequences.

## Acknowledgements

The computational work for this article was partially performed on resources of the National Supercomputing Centre (NSCC), Singapore (https://www.nscc.sg). JB is supported by the Singapore International Graduate Award (SINGA), A*STAR, Singapore. HF is supported by funding from Bioinformatics Institute, A*STAR, Singapore.

Appendix A. Supplementary Material

