## Supplementary Material for "A Molecular Dynamics Protocol for Rapid Prediction of EGFR Overactivation and Its Application to the Rare Mutations S768I, S768N, D761N"

### Appendix A. Supplementary Material

**Table S1**. Overview of simulated systems and simulation time in the benchmarking study.

| Force field | Variant | Dimer | Apo [ns] | Holo [ns] |
|---|---|---|---|---|
| CHARMM | WT | asymmetric | 500 | 500 |
| | | symmetric | 500 | 500 |
| | L858R | asymmetric | 500 | 500 |
| | | symmetric | 500 | 500 |
| | T790M/L858R | asymmetric | 500 | 500 |
| | | symmetric | 500 | 500 |
| AMBER | WT | asymmetric | 500 | 500 |
| | | symmetric | 500 | 500 |
| | L858R | asymmetric | 500 | 500 |
| | | symmetric | 500 | 500 |
| | T790M/L858R | asymmetric | 500 | 500 |
| | | symmetric | 500 | 500 |

**Table S2**. Overview of simulated systems and simulation time in the rare mutant study.

| Force field | Variant | Dimer | Apo [ns] | Holo [ns] |
|---|---|---|---|---|
| CHARMM | S768I | asymmetric | 500 | 500 |
| | | symmetric | 500 | 500 |
| | S768N | asymmetric | 500 | 500 |
| | | symmetric | 500 | 500 |
| | D761N | asymmetric | 500 | 500 |
| | | symmetric | 500 | 500 |

**Table S3**. Disruption of the inactive activation loop conformation in CHARMM simulations of the symmetric dimer. The activation loop was considered disrupted if the pitch of the short one-turn helix at its N-terminal end exceeded 7.0 Å. The table reports the percentage [%] of simulation time during which the loop was disrupted under Apo and Holo conditions.

| Mutant | Apo<br>Chain A | Apo<br>Chain B | Holo<br>Chain A | Holo<br>Chain B |
|---|---|---|---|---|
| WT | 38.7 | 12.1 | 15.3 | 11.8 |
| L858R | 70.6 | 80.6 | 93.8 | 52.5 |
| TMLR | 69.0 | 72.3 | 44.3 | 61.1 |
| S768I | 27.8 | 38.4 | 16.5 | 27.6 |
| S768N | 42.7 | 58.6 | 23.2 | 34.6 |
| D761N | 46.5 | 12.7 | 6.7 | 15.4 |

**Table S4**. Average number of hydrogen bonds formed by the side chain of residue 768 (Ser in WT, Asn in S768N) in CHARMM simulations of the symmetric dimer.

| Mutant | Apo<br>Chain A | Apo<br>Chain B | Holo<br>Chain A | Holo<br>Chain B |
|---|---|---|---|---|
| WT | 1.2 ± 0.5 | 1.1 ± 0.6 | 1.1 ± 0.6 | 1.2 ± 0.5 |
| S768N | 1.4 ± 1.2 | 1.8 ± 1.1 | 1.8 ± 1.2 | 1.7 ± 1.2 |
